# Tourism impacts on fecal glucocorticoid metabolites of free-ranging mountain gorillas in Bwindi Impenetrable National Park, Uganda

**DOI:** 10.64898/2026.01.15.699166

**Authors:** Ryoma Otsuka, Gladys Kalema-Zikusoka, Gen Yamakoshi, Kodzue Kinoshita

## Abstract

Mountain gorilla tourism plays a vital role in their conservation; however, increasing tourism intensity may elevate gorilla stress levels. Despite this concern, knowledge regarding its physiological effects remains limited. This study non-invasively assessed the impact of tourism on gorilla stress by measuring fecal glucocorticoid metabolites (fGCMs).

Fecal samples were collected from habituated gorilla groups in Bwindi Impenetrable National Park, Uganda, between 2017–2018. Concentrations of fGCMs were analyzed using a field-friendly extraction method and enzyme immunoassay. To assess the specific impacts of tourism, we focused on the Rushegura group, which is frequently visited by tourists, and used Bayesian linear mixed-effects models to examine associations between log-transformed fGCMs (log-fGCMs) and tourism intensity (daily tourist numbers and visit frequency). Additionally, we performed a inter-group comparison with the Mubare group, which was visited less frequently but underwent more dynamic social changes, to evaluate how log-fGCMs levels differed across different group contexts.

In the Rushegura group, the number of tourists and visits frequently exceeded the maximum limits established by the tourism guidelines (i.e., one visit of eight people per day). In this group, log-fGCMs increased with the number of tourists and were higher during days with multiple visits. The model incorporating visit frequency demonstrated better predictive performance. On the other hand, Mubare gorillas showed higher log-fGCMs than Rushegura gorillas, likely reflecting the influence of social instability.

These findings underscore the importance of adhering to the tourism guidelines—specifically regarding group size and visit frequency—to minimize physiological stress and associated risks, such as disease transmission. Although this pilot study lacks site-specific biological validation for this subspecies, the results provide preliminary insights into the physiological stress responses of Bwindi mountain gorillas to tourism. Further long-term, large-scale studies involving multiple gorilla groups are necessary to better understand how tourism and other factors (e.g., habituation status and social factors) influence their stress levels.

## Introduction

Mountain gorilla tourism is a very popular tourism activity in East Africa, and there is a growing number of tourists tracking gorillas. Mountain gorilla tourism has contributed to their conservation [1, 2] and the development of local communities [3]. However, tourism activity also poses risks to gorillas, such as disease transmission and/or stress [4]. There are tourism regulations/guidelines that aim to minimize such risks; however, there are pieces of evidence indicating the regulations/guidelines are not being strictly enforced or followed in the field [5–9]. Therefore, it is possible that the risk of tourism-related disease transmission and stress is increasing or may increase in the future if the situation remains unchanged.

While previous studies have provided important findings on disease transmission risks [10–12], the understanding of tourism impacts on physiological stress in mountain gorillas is very limited. A stress response is a natural, physiological, and behavioral coping mechanism to noxious stimuli; these are often called stressors [13]. When animals are confronted with a stressful situation, their adrenal cortex releases stress hormones such as cortisol or corticosterone into their blood (cf. [13–15]). The secretion of these hormones helps animals quickly respond to stressors through energy regulation [16]. Temporarily increased stress levels will be self-regulated through negative feedback in the hypothalamic-pituitary-adrenal axis [13]. However, if elevated stress levels persist for a long time and negative feedback is poorly regulated, this may have detrimental effects on the immune system and/or reproductive system of animals [16–18]. Given that there is a high risk of disease transmission between humans and gorillas [4, 10, 11], and that elevated stress levels can further increase susceptibility to illness [16, 17], the stress on gorillas associated with tourism should be minimized. Also, prolonged exposure to stressors, as seen in the habituation process, has been linked to increased parasitic loads, suggesting that chronic stress may exacerbate the health risks to gorillas [19].

In terms of the tourism impacts on behaviors of mountain gorillas, studies in Bwindi Impenetrable National Park, Uganda [9, 20] showed that the gorillas reduced the frequency of feeding and increased the frequency of moving, monitoring, self-directed behaviors, such as self-grooming and self-scratching, and human-directed behaviors when they were with tourists. Although self-grooming and self-scratching are often used as behavioral indicators of stress (or anxiety), an increase in these behaviors may not necessarily be linked to a rise in physiological stress levels [21]. Therefore, to better understand the physiological stress response of mountain gorillas to tourism activity, it is crucial to use non-invasive methods to measure more direct indicators of physiological stress.

Some studies non-invasively measured the physiological stress levels of mountain gorillas (through fecal or urine sample collection) [22–26]. However, these studies haven’t directly investigated the relationships between tourism and stress. In the western lowland gorilla population, Shutt et al. [27] investigated the impact of habituation and human activity (including tourism) on stress levels of gorillas in the Central African Republic, using a field-friendly, non-invasive method. The study showed that fecal glucocorticoid metabolites (fGCMs) levels were higher in the recently habituated group and the group undergoing habituation than in the non-habituated group, while there was no such difference between the fGCMs levels of the long-term habituated and those of unhabituated group, suggesting that fGCM levels return to baseline through long-term habituation [27]. While their analysis did not show an effect of the number of humans visiting the group, they further found that violation of the 7-m distance rule (i.e., tourists getting too close to gorillas) was associated with increased fGCMs levels in habituated gorillas. In other primate species, physiological stress responses to tourism vary across species and contexts [28–31], although it should be noted that these studies used different indicators of human (and/or tourism) activities. This underscores the importance of studying physiological stress in specific species or regions, as the unique traits of primate species and variations in tourism intensity (e.g., tourist density) and operational practices (e.g., proximity to animals) likely influence their stress responses to tourism.

In this study, we aimed to better understand the relationships between physiological stress levels and tourism activity in the Bwindi mountain gorilla population by monitoring fGCMs concentrations using a non-invasive method. Specifically, we examined the association between log-transformed fGCMs concentrations (log-fGCMs) and the intensity of tourism activity (the number of tourists or tourist visits per day) using Bayesian linear mixed-effects models with data from the Rushegura group, which was frequently visited by tourists during the fieldwork period. We also examined the effects of social (sex and age-class category) and ecological (mean temperature) factors that potentially influence the fGCMs levels of mountain gorillas [25–27]. To contextualize tourism’s impact, we compared the Rushegura group with the Mubare group, which experienced lower tourist visitation but underwent more dynamic social changes (i.e., a silverback takeover, female dispersal, and suspected infanticide). While not a direct comparison, this helps us gauge the magnitude of tourism-induced stress relative to that associated with natural social stressors.

## Materials and Methods

### Study area

We conducted fieldwork in Bwindi Impenetrable National Park, Uganda (0°53’–1°08’N; 29°35’–29°50’E; Figures 1 & 2a) which is characterized by many steep hills and narrow valleys [32]. The Bwindi forest covers approximately 331 km^2^ and the elevation ranges from 1,160 to 2,607 m above sea level [32]. The northern sector is the lowest sector in terms of altitude: its highest elevation is approximately 1,800 or 1,900 m above sea level. The annual mean minimum and maximum temperature ranges are 7–15°C and 20–27°C, respectively [33], while annual rainfall ranges between 1,400 and 1,900 mm [34]. There are usually two rainy (March-May and September–November) and dry seasons (December–February and June–August) within a year [35]. According to a survey conducted in 2018 in the Bwindi-Sarambwe region [35], a minimum of 459 mountain gorillas (approximately 43% of the total population of 1,063 [2, 35]) were estimated to inhabit the area.

**Figure 1.**
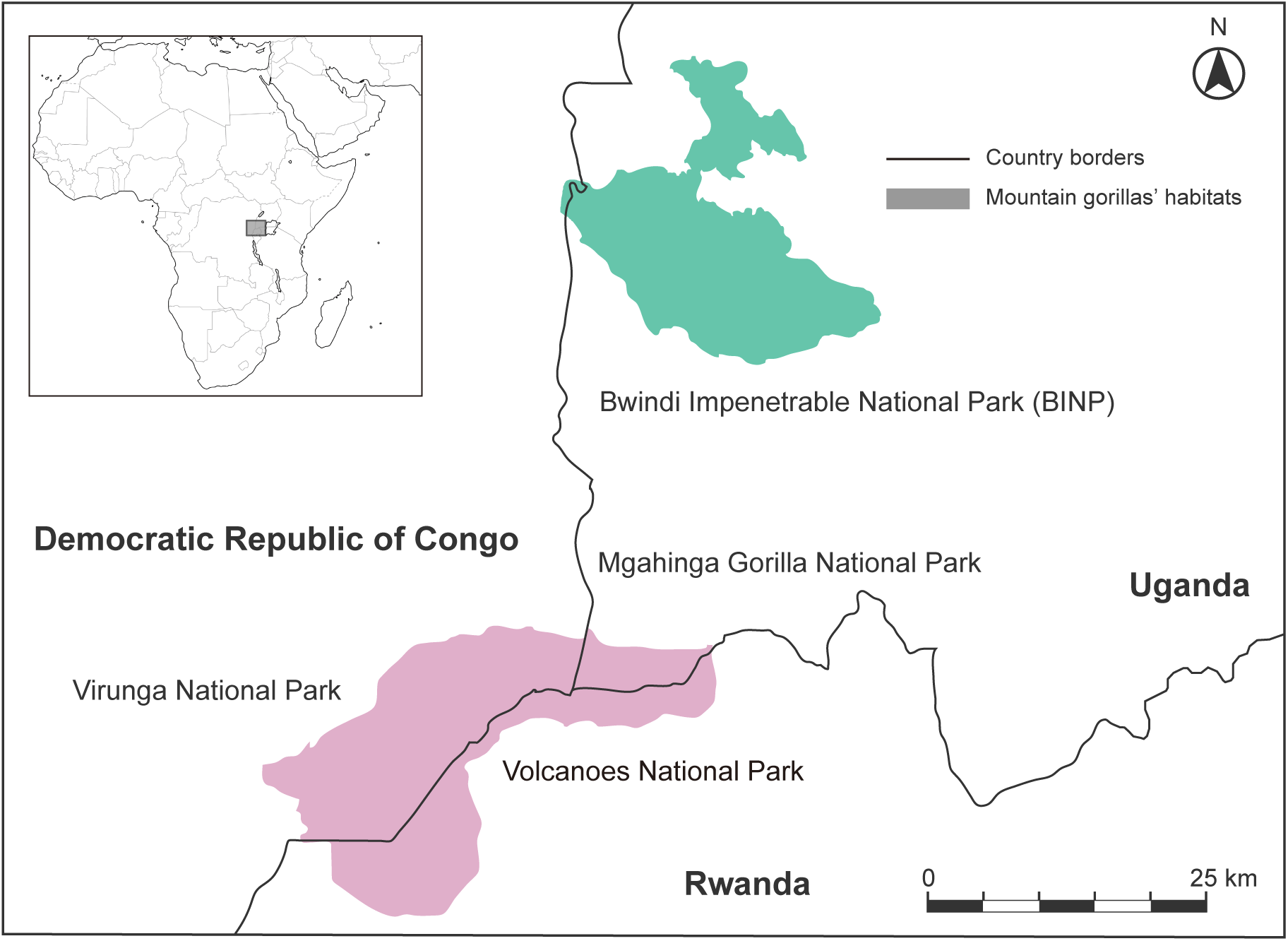
A map of Bwindi Impenetrable National Park, Uganda, and other mountain gorilla habitats.

**Figure 2.**
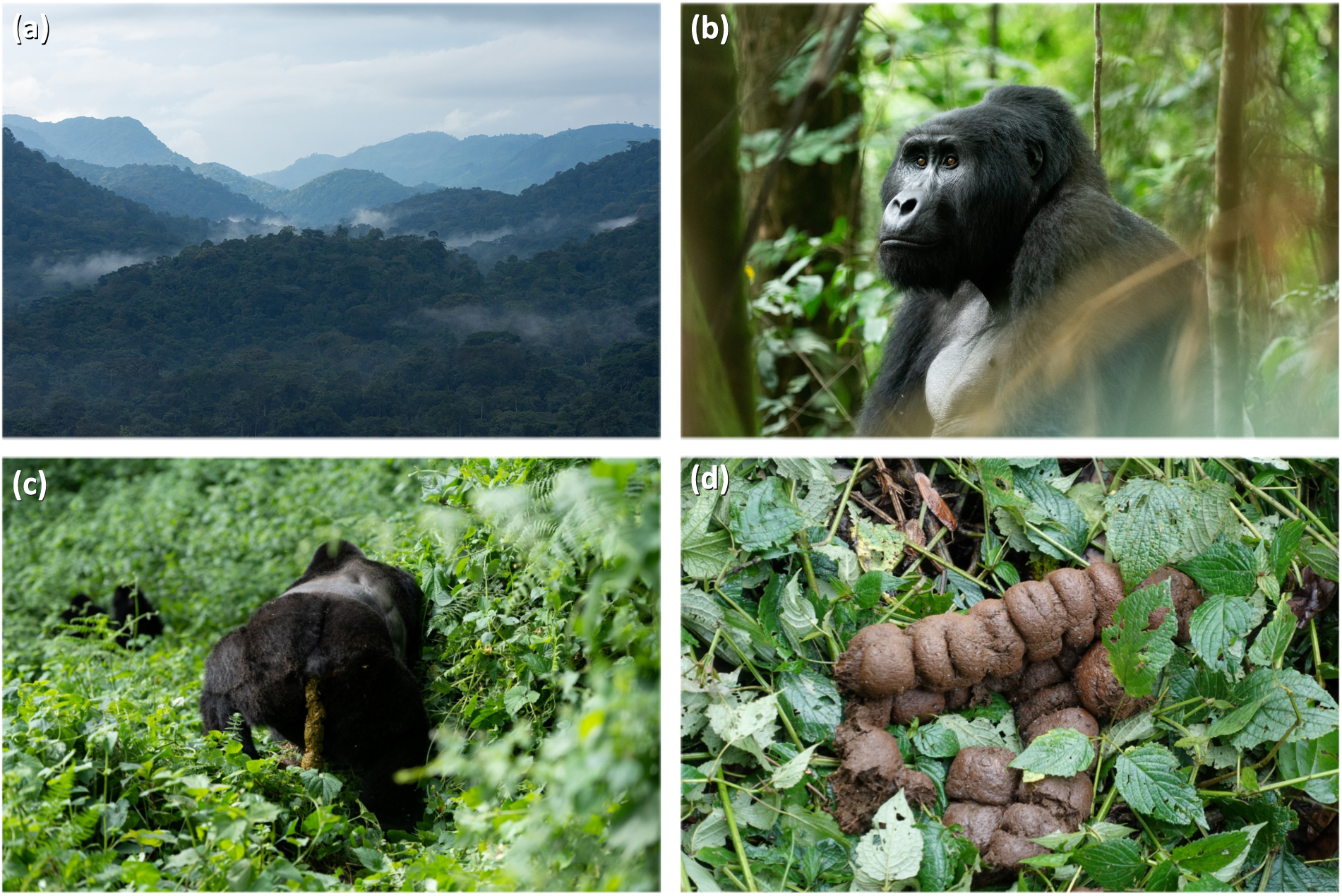
Photos of (a) Bwindi Impenetrable National Park; (b) Kanyonyi, the former silverback gorilla of the Mubare group; (c) a gorilla defecating; and (d) fecal samples.

Mountain gorilla tourism was introduced in Bwindi in conjunction with the Integrated Development and Conservation Projects (ICDP) [3, 36], beginning in 1993 in the northern sector [37]. The annual number of tourists visiting this park increased from 1,313 in 1993 to approximately 10,000 in 2008 [38]. Recently, the number of visitors is estimated to be approximately 40,000 each year [39], and most tourists visit the park to track and observe endangered mountain gorillas. The park is surrounded by high human population density (e.g., [40]), and a portion of the tourism revenue has been shared with communities, although there are also challenges in its implementation and system [38]. Regulations or guidelines for mountain gorilla tourism in Bwindi include a limited number of tourists (a maximum of eight tourists in one tourist group), a limited number of tourist visits, and regulated observation times (only one tourist group can track and observe one gorilla group in one day for a maximum of 1 h), and a 7 m distance rule [4].

### Study animals

There were four habituated groups of mountain gorillas in the northern sector of Bwindi during the fieldwork: the Habinyanja, Katwe, Mubare, and Rushegura groups. Among those, we mainly collected fecal samples from Rushegura and Mubare (Table 1 & 2). Very few samples could be collected from the Habinyanja group because they generally ranged far from our field camp (see also [41]), and the Katwe group was not yet well habituated, making tracking difficult. In fact, most samples collected from the Katwe group were nest samples, and we were unable to identify individuals from these samples.

**Table 1.** Detailed list of gorilla individuals and the number of fecal samples used in the model-R1 and model-R2.

| Gorilla ID | Group | Gorilla Name | Age Class | Sex | N of fecal samples |
| --- | --- | --- | --- | --- | --- |
| 1 | Rushegura | Barekye | Inf | M | 8 |
| 2 | Rushegura | Buzinza | AdF | F | 24 |
| 3 | Rushegura | Kabukojo | SB | M | 46 |
| 4 | Rushegura | Kabunga | SAd | M | 23 |
| 5 | Rushegura | Kankwehe | Inf | F | 7 |
| 6 | Rushegura | Kanyindo | AdF | F | 15 |
| 7 | Rushegura | Kanywani | BB | M | 23 |
| 8 | Rushegura | Karembezi | BB | M | 26 |
| 9 | Rushegura | Kibande | AdF | F | 32 |
| 10 | Rushegura | Masanyu | Inf | M | 10 |
| 11 | Rushegura | Mitunu | AdF | F | 25 |
| 12 | Rushegura | Munyana | AdF | F | 22 |
| 13 | Rushegura | Nyampazi | AdF | F | 24 |
| 14 | Rushegura | Rushokye | Inf | M | 6 |
| 15 | Rushegura | Ruterana | AdF | F | 29 |
| **Total** |  |  |  |  | **320** |

**Table 2.** Detailed list of gorilla individuals and the number of fecal samples used in the model-RM.

| Gorilla ID | Group | Gorilla Name | Age Class | Sex | N of fecal samples |
| --- | --- | --- | --- | --- | --- |
| 1 | Rushegura | Barekye | Inf | M | 8 |
| 2 | Rushegura | Buzinza | AdF | F | 25 |
| 3 | Rushegura | Kabukojo | SB | M | 51 |
| 4 | Rushegura | Kabunga | SAd | M | 23 |
| 5 | Rushegura | Kankwehe | Inf | F | 7 |
| 6 | Rushegura | Kanyindo | AdF | F | 18 |
| 7 | Rushegura | Kanywani | BB | M | 24 |
| 8 | Rushegura | Karembezi | BB | M | 26 |
| 9 | Rushegura | Kibande | AdF | F | 34 |
| 10 | Rushegura | Masanyu | Inf | M | 10 |
| 11 | Rushegura | Mitunu | AdF | F | 27 |
| 12 | Rushegura | Munyana | AdF | F | 26 |
| 13 | Rushegura | Nyampazi | AdF | F | 24 |
| 14 | Rushegura | Rushokye | Inf | M | 7 |
| 15 | Rushegura | Ruterana | AdF | F | 34 |
| 16 | Mubare | Karungi | AdF | F | 24 |
| 17 | Mubare | Kisho | AdF | F | 10 |
| 18 | Mubare | Kikombe | AdF | F | 15 |
| 19 | Mubare | Maraya | SB | M | 18 |
| 20 | Mubare | Twesiime | AdF | F | 14 |
| **Total** |  |  |  |  | **425** |

During our fieldwork period, the Mubare group experienced a drastic change in the social unit, and this event also affected the Rushegura group. The former silverback of the Mubare group, Kanyonyi (Figure 2b), died on 9 December 2017, after multiple attacks by a solitary silverback named Maraya (see [42] for more details). Until the death of Kanyonyi, the Mubare group consisted of one silverback, seven adult females, and six infants. Following his death, some group members dispersed, while some individuals remained and formed a social unit with the silverback, Maraya. This social unit continued to be called the Mubare group. During the fieldwork period in 2018, the Mubare group consisted of one silverback and four adult females (all were originally in the Mubare group), and zero infants.

On 1 June 2018, two adult females immigrated to the Rushegura group with their 3.25-year-old and 1.75-year-old infants; however, the younger infant later died because of infanticide [43]. These adult females were originally part of the Mubare group. Following these changes to the Rushegura group, the group consisted of a total of 15 members (one silverback, two blackbacks, seven adult females, one subadult, and four infants) up until the end of our study period. During the period after Maraya took over the leadership, the Rushegura group remained more tractable than the Mubare group, allowing for more efficient observation and sampling. The Rushegura group was also the most frequently visited group by tourists.

### Fecal sample collection, extraction, and preservation in the field

From August to October 2017, May to June, and September to October 2018, the first author and park rangers, who had been specifically trained in our sampling protocols, tracked gorillas and collected fresh fecal samples from trails in a non-invasive manner (Figure 2cd). Typically, gorilla tracking started at 07:00, and researchers were allowed to follow gorillas for up to 4 h per day, after the first sighting. We attempted to collect samples from as many individuals as possible while attempting to avoid imposing any additional stressors using various strategies, such as maintaining a sufficient distance from gorillas. We used small plastic bags (Unipack E-4 B7 100×140 mm, Seinichi) and laboratory gloves for sample collection. For trail samples, we collected fresh samples immediately after defecation; however, at times, there were waits of up to 30 min after defecation when gorillas were still around the dung. We recorded the date, time, and individual gorilla names for all samples collected. Gorilla individuals were identified by the first author, other researchers, and experienced guides and trackers. When an individual could not be identified, we did not collect samples from trails. We also did not collect samples when fecal samples were contaminated by urine. Samples were taken from a randomly selected section of the feces (proximal, middle, or distal) as fGCMs in gorilla dung are evenly distributed [23]. Regardless of the section chosen, we sampled only the interior portion of the fecal matter to prevent contamination from the environment. We also collected samples from nests, but the nest samples were used only for supplementary experiments. For nest samples, we recorded the dung diameter in addition to the date and time.

We transported samples to our camp and extracted hormones on the same day of collection. We placed fecal samples in an ice bag soon after the sample collection and tried to avoid direct sunshine until returning to the camp. The samples were refrigerated at 4°C immediately after returning to the camp. We extracted hormones approximately 1 to 2 h after returning to the camp; hormone extractions typically occurred 2 to 10 h after sample collection. For hormone extraction and preservation, we followed a field-friendly method that was proposed and validated by Shutt et al. [44]. This is mainly because sufficient electric power for the fridge to maintain the temperature below −20°C and other experimental equipments were not available. Prior to hormone extraction, each sample was manually homogenized in a plastic bag. Then, 1.00 g of fecal matter was removed from each sample and weighed using a portable electronic scale (Personal Electronic Scale EJ-120, A & D), and immediately placed into a 25 mL self-standing bottle (Centrifuge Tube Mini, IWAKI) containing 8.0 mL of 90% ethanol. Bottles were manually shaken horizontally for 5 min, and then left undisturbed on a laboratory bench for 40 min. After 40 min, 0.8 mL of supernatant was transferred into a 1.5 mL microtube (1.5 mL Flat-Bottom Micro Tube, WATSON). These were dried at room temperature in the dark until they were completely dried. For drying process, we used silica gel because of the very high humidity in Bwindi. After drying, we closed the cap of each tube and stored these tubes at room temperature in the dark until shipment.

We have conducted a series of field experiments (Experiments S1–4) to better understand the pros and cons of various field sampling and extraction techniques. See supplementary experiments and Figures S1 & S2 in the supplementary information for more details.

The research permits were provided by the Uganda Wildlife Authority (UWA) and the Uganda National Council for Science and Technology (UNCST). For sample collection and transfer, a Material Transfer Agreement (MTA) was obtained from UWA, and an export permit for the samples was provided by the UNCST.

### Measurement of fGCMs concentrations using enzyme immunoassay

All dried samples were transported to Japan and stored in a freezer (−20°C) at the Wildlife Research Center (WRC), Kyoto University, until measurement. We measured fGCMs concentration using EIA at the WRC laboratory between January and March 2019. We followed the protocol for cortisol EIA with the FKA404E antibody and FKA403 antigen (Cosmo Bio Co., Ltd.), which has been used for other mammal species, including non-human primates [31, 45]. While a species-specific validation (e.g., ACTH challenge or biological validation) is ideally required, such invasive procedures and sufficient behavioral/physiological data for biological validation were not feasible in this study. Although this remains a primary limitation of the present work, the use of this well-established antibody/antigen system provides a plausible proxy for cortisol measurement in this species. Dried samples were reconstituted using 0.8 mL of 80% methanol.

Sample extracts were diluted by a factor of 15 and assayed in duplicate. The absorbance of each well in each plate was measured at 450 nm using a microplate reader (SUNRISE, BIO RAD Laboratories Inc.). Measured fGCMs concentrations were corrected using synthetic quality control samples included on each plate. In total, we measured the fGCMs concentrations of 965 samples including samples for a series of experiments (Experiments S1–4). The inter-assay (29 plates) and intra-assay (mean coefficient of variation of duplicate measurements for each sample within a plate) coefficients of variation (CV) were 18.83% and 5.27%, respectively. The parallelism of the assay was evaluated by comparing serially diluted samples with the cortisol standard curve. We visually confirmed that the slopes of the samples and the standard curve were nearly parallel (Figure S3).

### Data analysis

#### Model structure

Data analysis was conducted using R 4.1.3 [46]. Data and source code are available from the GitHub repository (https://github.com/ryoma-otsuka/mg-fgcms). We constructed and fitted three different Bayesian linear mixed-effects models: model-R1, model-R2, and model-RM, where “R” and “M” denote the Rushegura and Mubare groups, respectively. We used the model-R1 and model-R2 to explore the impact of tourism activity on the physiological stress levels of gorillas in the Rushegura group. On the other hand, we used model-RM to compare the Rushegura group, which experienced substantially higher tourism pressure, with the Mubare group, which was less frequently exposed to tourism but experienced social instability.

Although not a direct comparison, this allowed us to contextualize the magnitude of tourism-induced physiological responses relative to those triggered by natural social stressors. In this section, we describe the common model structures and settings for model fitting. The basic structure of these models is common, but the explanatory variables and dataset are different. In the following sections, we further describe the details of the explanatory variables and dataset used in these models. The structure of the Bayesian linear mixed-effects models is defined as follows:

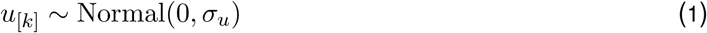

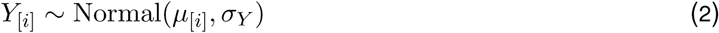

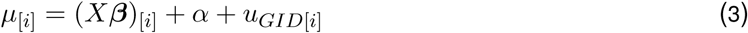

As the response variable *Y*_[*i*]_, we used log-transformed fGCM concentrations (hereafter, log-fGCMs), following previous studies [26, 27]. We checked the distribution of the raw fGCMs concentration data (Figure S4) and assumed that there might be exponential relationships between the response variable and explanatory variables. The parameter *α* represents the global intercept. To account for individual-specific baselines in fGCM levels, we included gorilla individual identification (GID) as a random intercept *u_GID[k]_*, where *σ_u_* denotes the standard deviation between individuals. The term *X****β*** represents the fixed effects component, where *X* is the design matrix of explanatory variables (e.g., number of tourists, visit frequency) and ***β*** is a vector of the corresponding slope parameters (or regression coefficients) to be estimated. Finally, *σ_Y_* represents the standard deviation, reflecting the unexplained variation in the log-fGCM data. For all models, we employed weakly informative priors. The specific prior distributions were defined as follows:

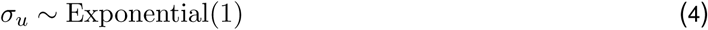

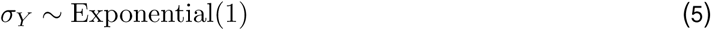

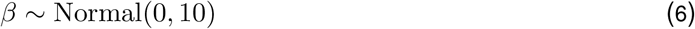

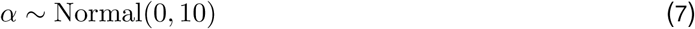

We used Stan 2.21.0 [47] and RStan 2.21.7 [48] to run Markov chain Monte Carlo (MCMC) sampling. For MCMC sampling, we used four chains, 4,000 iterations with 2,000 warm-up, and samples were not thinned out. We set the adapt delta to 0.99. We checked the convergence by visually examining trace plots (e.g., we checked if each chain was well mixed in trace plots). We also confirmed that the Gelman-Rubin statistic (the Rhat value) for each parameter was less than 1.01. To verify if the models appropriately regenerated the data, posterior predictive checks were conducted by drawing randomly extracted 100 posterior predictive density distributions from each model along with the probability density distribution of observed data. The posterior predictive checks confirmed that the models adequately regenerated the observations (Figures 3 & 6a). We show the posterior median and 89% and 97% Bayesian credible interval (BCI) of the posterior distributions in the results.

**Figure 3.**
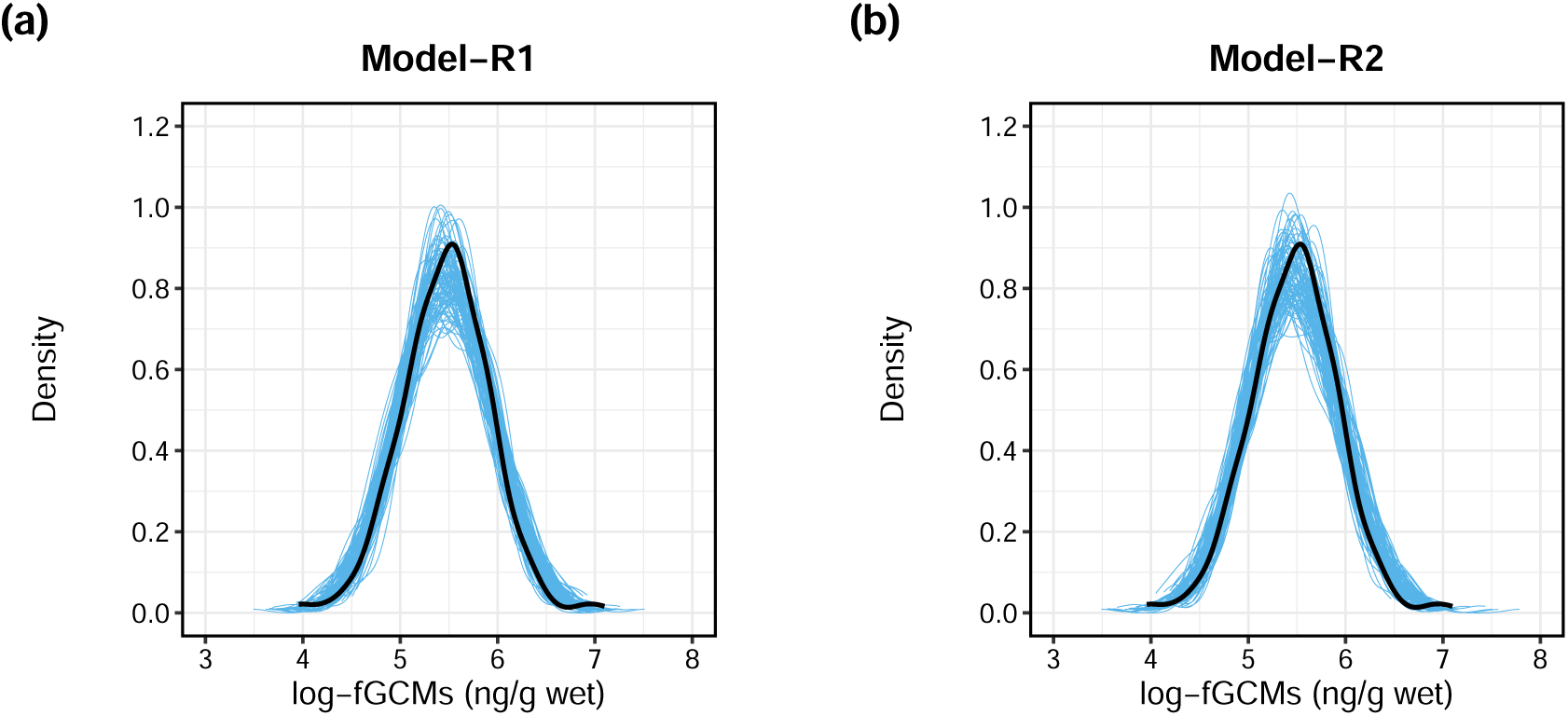
Posterior predictive checks for the model-R1 (a) and the model-R2 (b). The blue lines are the randomly extracted 100 posterior predictive probability density distributions, while the black line shows the probability density distribution of observed data.

#### Explanatory variables for Model-R1 and Model-R2

The model-R1 and model-R2 included common explanatory variables: sex, age-category, and daily mean temperature. As socio-demographic factors, we included gorilla sex (female/male; coded as a dummy variable where 0 = female and 1 = male) and gorilla age category (adult/infant; coded as a dummy variable where 0 = adult and 1 = infant) in these models. We did not include age data as a numeric variable because the exact age of some gorillas was unknown. We generally followed the age classifications defined by Williamson and Gerald-Steklis [49], which categorize individuals over 8.0 years old as adults and those aged 0.0–3.5 years as infants. However, for the purpose of our analysis, two exceptions were made for the Rushegura group in this study. First, one individual aged approximately 7 years was treated as an adult. Second, another Rushegura individual, whose age slightly exceeded 3.5 years by the end of our fieldwork in October 2018, was classified as an infant.

As an environmental indicator, the daily mean temperature was also incorporated in the models. The temperature was measured every hour using a digital thermometer and hygrometer (Thermo Recorder Ondotori TR72-wf-H, T & D), installed at the Buhoma field station of the UWA at 1.5 m above the ground surface. We standardized the daily mean temperature by subtracting the mean value and dividing it by the standard deviation prior to fitting the models. Unfortunately, rainfall data were not available for use in this study.

As indicators of tourism activity, we used the daily total number of tourists and the daily total number of tourist visits. Specifically, the daily total number of tourists refers to the aggregate number of visitors per group per day, including all individuals across multiple visiting sessions. These data were collected primarily by field observations. To ensure maximum data accuracy and completeness, we cross-referenced our records with data provided by UWA and other researchers, consolidating multiple sources to refine the final dataset. For tourism intensity indicators, model-R1 included the daily number of tourists as an indicator of tourism activity, whereas model-R2 included the daily number of tourist visits. As the daily total number of tourists and the daily total number of tourist visits were highly correlated, we analysed the impacts of these variables in the different models. We standardized the daily total number of tourists by subtracting the mean value and dividing it by the standard deviation prior to fitting the models. The daily number of tourist visits was included as a categorical variable (once/twice/three times; coded as dummy variables where 0 = once, 1 = twice, and 2 = three times) in the model-R2. Unfortunately, behavioral data of gorillas and/or tourists were not incorporated into the model, as we were unable to collect enough behavioral data along with the fecal sample collection.

Shutt et al. [27] used a 48-h time lag because they previously found fGCMs concentrations peak 48 h after a stressor in western lowland gorillas [44]. Nizeyi et al. [24] revealed that fGCMs concentrations were elevated between 72 and 96 h (if measured by RIA), and between 48 and 120 h (if measured by EIA) following the injection of the adrenocorticotrophic hormone to captive western lowland gorillas. They also found that fGCMs concentrations increased between the second and third day of gorillas being chased by local people from crop fields for three free-ranging mountain gorillas in Bwindi, although fGCMs concentrations were measured using RIA in this field validation [24]. In the biological validation for the Virunga mountain gorilla population, [25] showed that fGCMs concentrations increased between 20 and 140 h after inter-unit interactions, and the peak often occurred on the third day after interactions. It is known that Bwindi mountain gorillas reside at lower altitudes and are more frugivorous than Virunga mountain gorillas (Robbins and McNeilage 2003; Ganas et al. 2004; Nkurunungi et al. 2004). The gut passage time may be affected by this unique feeding ecology and generally, the gut passage time is expected to be shorter in frugivorous species than in herbivorous species [50]. Therefore, we used a 48-h time lag in this study. For the daily mean temperature, daily total number of tourists, and daily number of tourist visits, we used data measured two days prior to the date of fecal sample collection; this means that if a fecal sample was collected on 24 September 2018, we used data from 22 September 2018.

#### Model comparison of model-R1 and model-R2

Models-R1 and model-R2 differ in their choice of tourism indicator: model-R1 uses the daily total number of tourists, while model-R2 uses the daily total number of tourist visits. By comparing the out-of-sample predictive performance of these two models, we aimed to determine which indicator serves as a better predictor of gorilla stress levels. The out-of-sample prediction accuracy of the model-R1 and model-R2 were compared using the Widely Applicable Information Criterion (WAIC) [51] and Pareto Smoothed Importance Sampling Leave-One-Out Cross-Validation (PSIS-LOOCV) [52]. These metics were calculated and compared using the “compare” function in the “rethinking” package version 2.42 [53].

#### Explanatory variables for model-RM

The model-RM was used to investigate the differences in fGCMs levels between the Mubare and Rushegura groups. The model included sex, age-category, and group (Mubare/Rushegura; coded as a dummy variable where 0 = Mubare and 1 = Rushegura) as explanatory variables. The basic model structure, priors, and settings for MCMC sampling were the same as those of the model-R1 and model-R2.

#### Dataset for modelling

By examining the histogram of the raw data, we considered one sample from an adult female gorilla in the Mubare group (that was more than 8,000 ng/g) as an extreme observation, likely due to measurement error or contamination; therefore, this sample was excluded (see Figure S4). We removed observations that contained missing values in any of the explanatory variables. Ultimately, all samples collected in 2017 were excluded because of a lack of sufficient information and low sampling frequency. Only trail samples were used, and all nest samples were excluded from the analysis.

For the model-R1 and model-R2, we exclusively used the data from the Rushegura group. The Rushegura group visited by one or more tourist groups on a daily basis, providing a robust and consistent dataset for analysis. In contrast, the sample size obtained from the Mubare group was considerably smaller. Furthermore, during fieldwork in 2018, there were days in which the Mubare group received no tourist visits, likely reflecting the group’s instability at the time. The impacts of drastic changes in this group would make it difficult to simply analyze the relationship between fGCMs concentrations and tourism activity. Therefore, the Mubare group was excluded from these specific models to maintain analytical consistency. Consequently, a total of 320 samples from the Rushegura group were included in the analysis. These samples represent 15 distinct individuals; the specific breakdown of fecal samples per individual is detailed in Table 1.

For the model-RM, we utlized the data from the Rushegura and Mubare group. Samples collected in 2017 were excluded from this analysis for both groups to account for their limited sample sizes and dynamic changes in group composition, specifically the silverback takeover and subsequent immigrations and emigrations. Consequently, the dataset in this analysis contained 81 samples from the Mubare group and 344 samples from the Rushegura group (425 samples in total). The number of fecal samples for each individual gorilla (representing 20 individuals in total) is detailed in Table 2.

## Results

### Model-R1 and Model-R2

The posterior distribution of *β*_4_ in the model-R1 indicated that there was a positive association between the daily total number of tourists and log-fGCMs (Figures 4a & 5de). The posterior distributions of *β*_4_ and *β*_5_ in the model-R2 indicated that log-fGCMs were higher when a gorilla group was visited by tourists twice or three times a day, compared to when a gorilla group was visited only once (Figures 4b & 5f). Although the wide uncertainty of *β*_5_ in the model-R2 may be due to the small sample size for the category (i.e., number of tourist visits = 3, n = 15), these results suggest that gorillas experienced higher stress levels as both the total number of tourists and the frequency of separate tourist visits increased.

**Figure 4.**
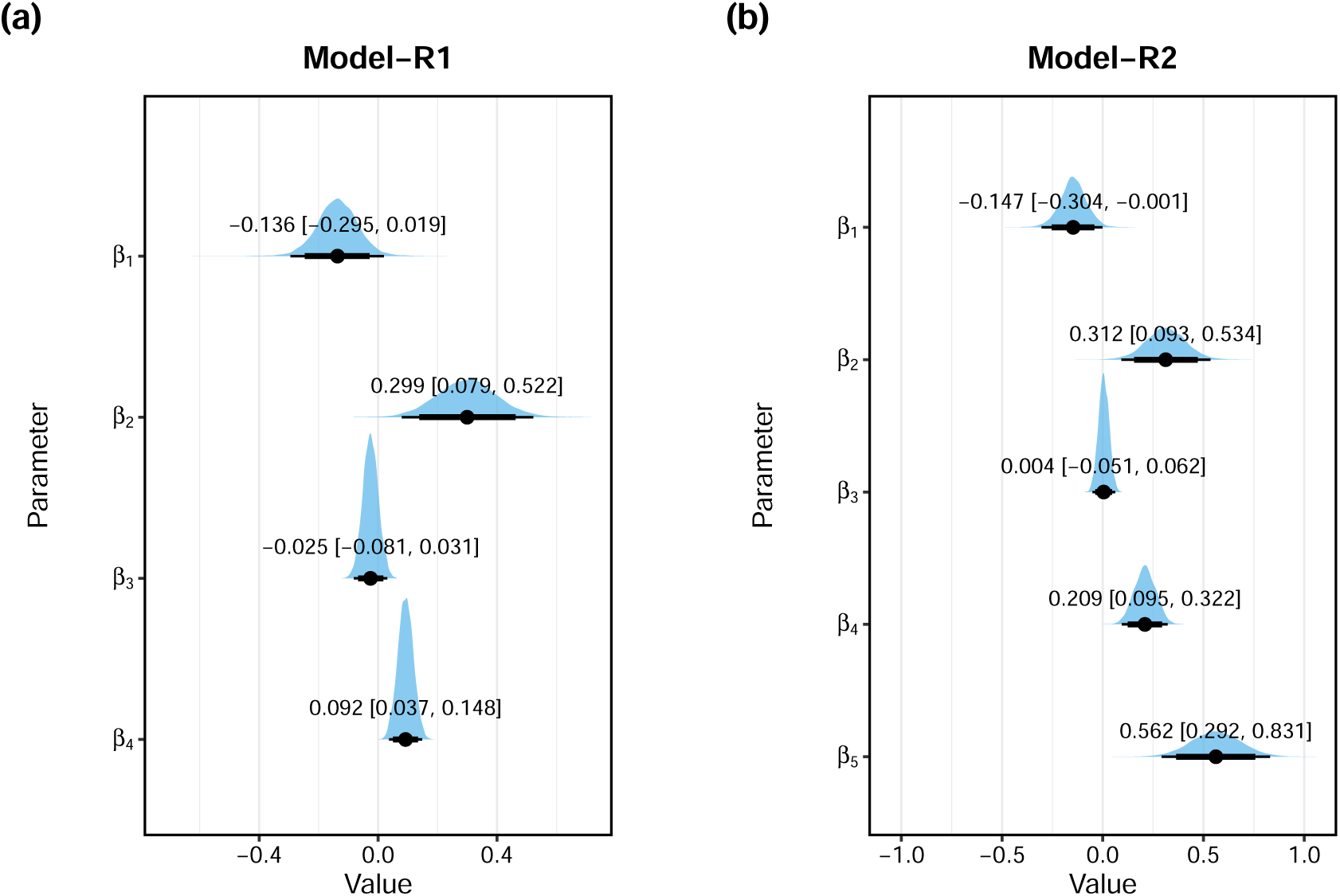
Posterior distributions of slope parameters from the model-R1 (a) and the model-R2 (b). The blue half-eye-shaped objects represent posterior distributions. The black circles indicate the median, while thick and thin black lines indicate 89% and 97% Bayesian credible intervals (BCIs), respectively. For the model-R1, the parameters *β*_1_–*β*_4_ are the slope parameters of the following variables: sex (0: female, 1: male), age-category (0: adult, 1: infant), the daily mean temperature, and the daily total number of tourists, respectively. For the model-R2, the parameters *β*_1_–*β*_5_ are the slope parameters of the following variables: sex (0: female, 1: male), age-category (0: adult, 1: infant), the daily mean temperature, and the daily total number of tourist visits for the group (= twice, three times), respectively. The posterior median values and 97% BCIs are shown as texts.

**Figure 5.**
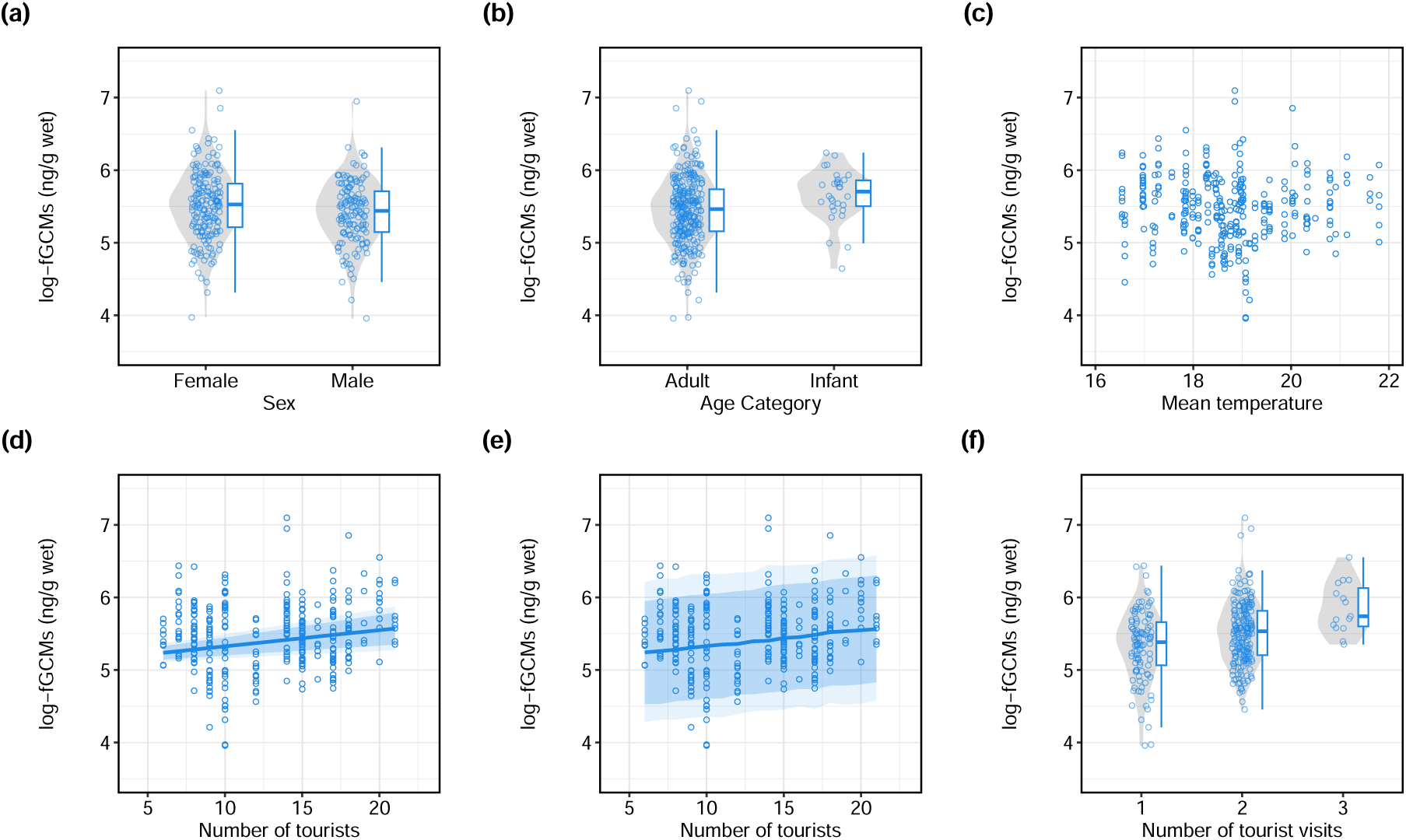
Relationships between log-transformed fGCMs concentrations and explanatory variables: sex (a), age-category (b), the daily mean temperature (c), the daily number of tourists (d, e), and the number of tourist visits (f) in the Rushegura group (N = 320). In (d), the regression line (median) and 89% and 97% Bayesian confidence intervals (BCIs) are shown. In (e), the prediction line (median) and 89% and 97% Bayesian prediction intervals are shown. To visualize median lines and BCIs, we used the posterior median of parameter *β*_3_ of the model-R1 and the mean value for daily mean temperature, while fixing gorilla sex as female (sex = 0) and age-category as adult (age-category = 0).

Model comparisons showed that the WAIC and PSIS values of the model-R2 were smaller than those of the model-R1 by approximately 13.0 (Tables 3 & 4). This indicates that the model-R2 outperforms the model-R1 in terms of out-of-sample prediction accuracy. Given that the only difference between the two models was a tourism-related variable used (i.e., the daily total number of tourists or the daily number of tourist visits), the difference in prediction accuracy originates from the these respective variables. This suggests that repeated exposure to tourist groups can be a more critical determinant of physiological stress than the total number of visitors.

Both models showed similar results for the slope parameters (i.e., *β*_1_ – *β*_3_) of the common explanatory variables. The log-fGCMs of male gorillas were slightly lower than those of female gorillas (*β*_1_ in Figures 4 & 5a). Infant gorillas had relatively higher log-fGCMs than adult gorillas (*β*_2_ in Figures 4 & 5b). The wide uncertainty of this parameter may be due to the small sample size for the infant category (n = 30). There was no clear association between the daily mean temperature and log-fGCMs in either model (Figures 4 & 5c).

### Model-RM

The posterior distributions of *β*_1_ and *β*_2_ from the model-RM showed similar trends with those from the model-R1 and model-R2, and the posterior distribution of *β*_3_ indicated that gorillas in the Rushegura group had lower log-fGCMs than those in the Mubare group (Figures 6b & 7).

**Figure 6.**
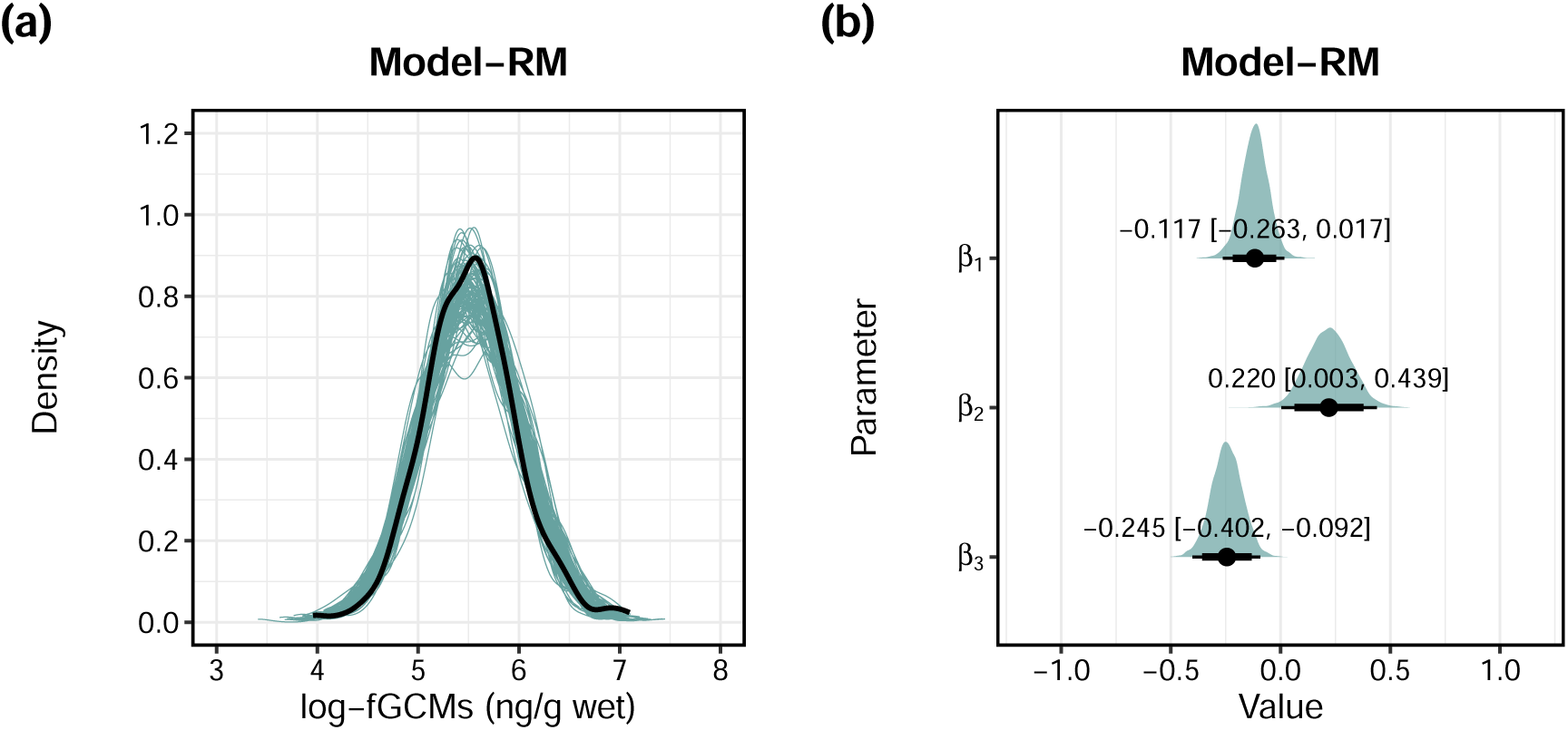
Posterior predictive check for model-RM (a) and posterior distributions of slope parameters (b). For (a), the green lines are the randomly extracted 100 posterior predictive probability density distributions, while the black line shows the probability density distribution of observed data. For (b), the green half-eye-shaped objects represent posterior distributions. The black circles indicate the median, while thick and thin black lines indicate 89% and 97% Bayesian credible intervals (BCI), respectively. The parameters *β*_1_–*β*_3_ are the slope parameters of the following variables: sex (0: female, 1: male), age-category (0: adult, 1: infant), and group (0: Mubare, 1: Rushegura), respectively.

**Figure 7.**
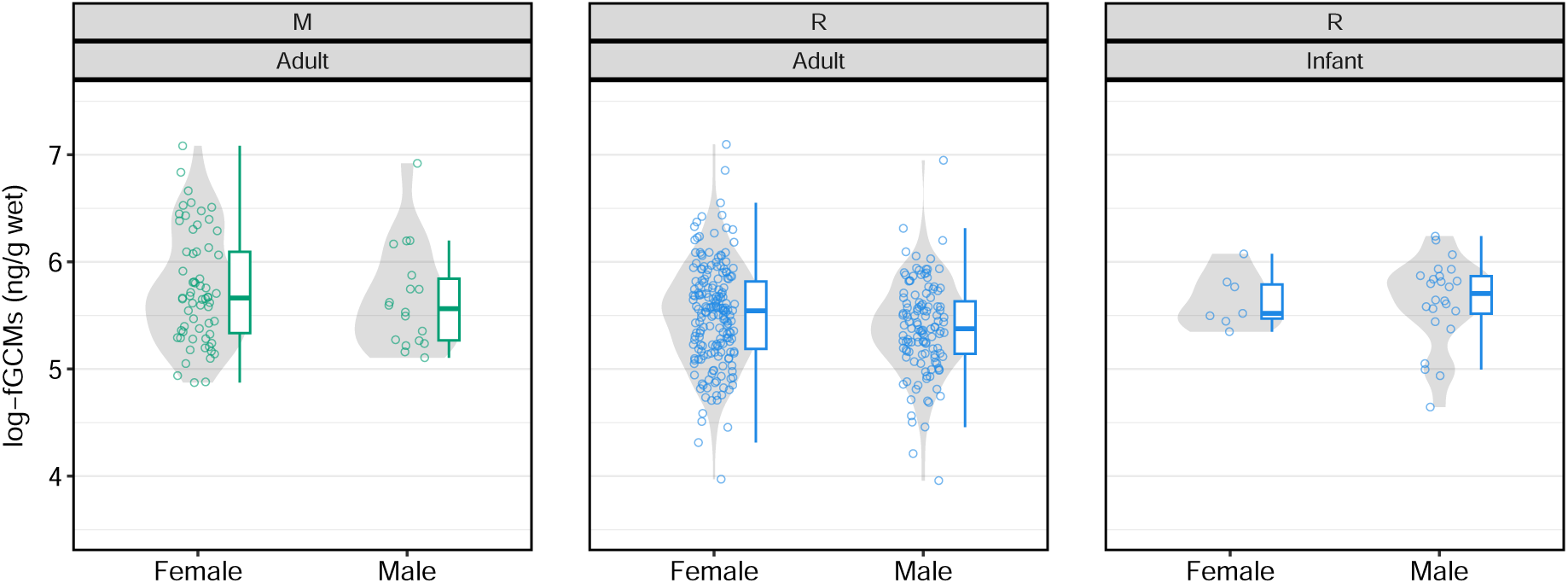
Comparison of log-transformed fGCMs concentrations of samples from gorillas in the Mubare group (N = 81, 63 samples from females and 18 samples from a male, no infant was in this group), and the Rushegura group (N = 344, 196 samples from females and 148 samples from males, 313 samples were from adults and 31 samples were from infants).

**Table 3.** Model comparison using WAIC.

| Model | WAIC | SE | dWAIC | dSE | pWAIC | weight |
| --- | --- | --- | --- | --- | --- | --- |
| Model-R2 | 383.2139 | 30.3379 | 0 | NA | 10.3689 | 0.9985 |
| Model-R1 | 396.2290 | 30.4711 | 13.0151 | 5.4074 | 9.5644 | 0.0015 |

**Table 4.** Model comparison using PSIS-LOOCV.

| Model | PSIS | SE | dPSIS | dSE | pPSIS | weight |
| --- | --- | --- | --- | --- | --- | --- |
| Model-R2 | 383.3141 | 30.3978 | 0.0000 | NA | 10.4190 | 0.9985 |
| Model-R1 | 396.3150 | 30.5313 | 13.0008 | 5.4070 | 9.6074 | 0.0015 |

## Discussion

### Tourism impacts

To our knowledge, this study is the first to report on the relationships between tourism activity and fGCMs concentrations in the mountain gorilla population. Our findings demonstrate that both the daily total number of tourists and the daily number of tourist visits were linked to elevated physiological stress levels in the Rushegura gorillas. Our results show that stress levels tended to rise with both higher aggregate visitor numbers and increased visit frequency, with the latter serving as a more decisive predictor of the stress response.

The number of tourists was not associated with fGCMs concentrations in Japanese macaques in Jigokudani, Japan [31] and Barbary macaques in Morocco [29]. Further, Shutt et al. [27] showed that the number of humans, such as tourists and researchers, who encountered the studied gorilla groups, was not associated with fGCMs concentrations in the western gorillas in Bai Hokou and Mongambe. While the number of tourists was restricted to a maximum of six people (including a guide and two trackers) per group in Bai Hokou and Mongambe [27], a maximum number of eight tourists per day per group were permitted in the mountain gorilla field sites [4]. In addition, during the study period, the studied group was sometimes visited by more than eight tourists. Thus, it is likely that there were fewer variations in terms of the daily number of tourists (or humans, including researchers and rangers) for western lowland gorillas in Bai Hokou and Mongambe than the mountain gorilla population. The model-R1 suggested that the total number of tourists per day was positively associated with log-fGCMs concentrations. However, the total number of tourists normally increases with the number of tourist visits. Model comparison using WAIC and PSIS-LOOCV revealed that the model-R2 was better in terms of out-of-sample prediction accuracy. In other words, the results suggest that the model-R2, which included the daily number of tourist visits as a tourism indicator, better predicts the tourism impacts on the stress response in free-ranging mountain gorillas in Bwindi.

The studied groups were often visited twice (or three times) a day by different groups of tourists during the study period. It is unclear whether this phenomenon was only observed during the study period or if it has been common. This may be because the newly habituated group, the Katwe, was unavailable. It is also possible that a well-habituated group like the Rushegura group, containing an infant, was considered to be more attractive for tourists than a group without any infant, like the Mubare group, during the study period in 2018. A difference in tracking difficulty of each group may also be the cause of this phenomenon. In fact, the Rushegura group had been relatively easier to track than other groups. The study in the western lowland gorilla population observed an approximately 50% increase in fGCMs concentrations 48 h after the gorillas in the group undergoing habituation encountered humans humans three times in one day, while this did not occur when they were only encountered once or twice a day [27]. It is possible that an increased number of contacts with humans within one day is associated with increased fGCMs concentrations of free-ranging gorillas. This may be attributed to the increased costs of moving and monitoring, and/or the reduced feeding time during tourist visits [9, 20]. As one visit by one tourist group lasts for approximately 1 h [6], this means that gorillas spend 2–3 h under observation by tourists when visited twice or three times a day. It is reasonable that repeated encounters with unfamiliar humans (being approached by strangers) several times a day may elevate the stress levels of Bwindi mountain gorillas.

Previous studies highlight the importance of habituation levels. Shutt et al. [27] demonstrated that the fGCMs concentrations of gorillas in human-contacted groups (particularly recently habituated groups and groups undergoing habituation) were higher than those of gorillas in the unhabituated group. However, they did not find such a difference between long-term habituated and unhabituated groups. A study on habituated orangutans concluded that habituated orangutans were not chronically stressed by tourism [30]. They found that, although stress levels increased after contact with tourists, the stress levels of habituated orangutans were lower than their age-matched unhabituated orangutans [30]. The present study did not examine whether habituated Bwindi mountain gorillas are at a higher risk of chronic stress than unhabituated gorillas. Therefore, incorporating groups across a broader spectrum of habituation levels in future research would be highly beneficial. Such long-term, large-scale comparisons are needed to clarify how habituation status and tourism activity together influence the chronic stress levels of mountain gorillas.

### Implications from the comparison between the two groups

Model-RM showed that the gorillas in the Mubare group had higher fGCMs concentrations than those in the Rushegura group. The Mubare group was less frequently visited by tourists than the Rushegura group. However, the Mubare group had recently experienced dynamic changes within the group, including the death of silverback gorillas, a takeover by a new silverback, the emigration of adult females, and suspected infanticide. This instability within the group may have contributed to the relatively higher fGCMs concentrations in the Mubare group than the Rushegura group. These observations suggest that while tourism influences fGCM levels, such increases may remain within the range associated with “natural” social stressors. Our results, therefore, highlight that the impact of tourism should be interpreted in conjunction with natural social dynamics. This also underscores the importance of long-term and large-scale studies to disentangle the relative contributions of anthropogenic and natural stressors on gorilla stress.

### Socio-demographic factors

Our results indicated that male gorillas had lower log-fGCMs concentrations than female gorillas, which is consistent with the previous studies on the Virunga mountain gorilla population [25, 26]. This is also consistent with the results of a study on captive western lowland gorillas [54], while the previous study in Bai Hokou and Mongambe [27] did not find a significant relationship between mean fGCMs concentrations and sex in western lowland gorillas. The difference in fGCMs concentrations between sexes found in this study may represent differences between sexes in terms of hormone metabolism [55]. Also, it may be related to other factors such as energy demands and female reproductive status, as discussed in Eckardt et al. [25].

In addition, the results indicated higher log-fGCMs concentrations in infant gorillas than in adult gorillas. While the study in Bai Hokou and Mongambe did not detect a difference between age-class [27], our result is consistent with the study in Virunga mountain gorillas, which demonstrated that elder gorillas had relatively lower baseline fGCMs concentrations than younger individuals [26]. Another previous study in Virunga mountain gorillas used urine samples and showed that immature males in Virunga had higher levels of cortisol than maturing and adult males [22]. It is known that age can affect the physiological stress levels in other mammal species (e.g., [56]) and is generally regarded as a potential factor that affects physiological stress levels [15, 57]. Overall, our findings on socio-demographic factors are consistent with general knowledge and support the results in the Virunga mountain gorilla population.

### Environmental factors

Eckardt and colleagues [26] demonstrated that the daily maximum and minimum temperature and rainfall were positively associated with higher baselines of fGCMs concentrations in the Virunga mountain gorilla population. Shutt et al. [27] did not find a significant relationship between mean fGCMs concentrations and mean daily temperature in western lowland gorillas in Bai Hokou and Mongambe. The models in this study showed no clear association between the daily mean temperature and log-fGCMs concentrations, which is consistent with the findings of Shutt et al. [27]. This may be because Bwindi mountain gorillas (our study groups in the northern sector of Bwindi, in particular) range at relatively lower altitudes compared to Virunga mountain gorillas. While our study groups ranged from approximately 1,450–1,800 m above sea level [58], the Virunga mountain gorillas studied in the previous research ranged from approximately 2,300–4,500 m [26]. At these lower altitudes, the minimum temperatures remain relatively mild compared to the high-elevation habitats in the Virunga. Consequently, the gorillas in our study area are less likely to experience severe cold stress or the significant thermoregulatory. However, it should be noted that we only used the mean temperature as an environmental factor. The impacts of other environmental factors, such as maximum and minimum temperature, humidity, and rainfall, on fGCMs concentrations should be investigated in future research for the Bwindi mountain gorilla population.

### Limitations

Limitations of this study include the small sample size, low frequency of sampling from each gorilla individual, short sampling period, lack of detailed observational data on gorilla behaviors, and the lack of biological validation for the specific sampling and preservation methods used for the Bwindi population. The main results of this study on tourism impacts are based on the data from only one gorilla group (i.e., the Rushegura group). To collect both fecal samples and behavior data from multiple groups, a large budget and sufficient manpower are needed. Unfortunately, we were unable to do that due to limited budget and manpower. In addition, detailed observational data on interactions within the studied gorilla group or interactions with other gorilla groups may explain changes in fGCMs concentrations. In this study, we did not consider such factors (e.g., inter-group interactions) that are known to have an influence on fGCMs levels [25, 26], due to a lack of data. The influence of interactions between gorillas and ranger guides, trackers, and local people should also be considered in future research. Finally, although the similar field-friendly methods applied in this study were biologically validated for the free-ranging western lowland gorilla population [44], the Bwindi mountain gorilla population [24], and the Virunga mountain gorilla population [25], the specific method used in this study was not biologically validated for Bwindi mountain gorillas. Future studies should carefully consider these limitations before starting the fieldwork.

## Conclusion

This study contributes to an understanding of the stress response to tourism activity by free-ranging mountain gorillas in Bwindi and suggests that the increased tourism activity may elevate the stress levels of mountain gorillas in Bwindi. Approximately 57% of the mountain gorilla population in the Bwindi-Sarambwe ecosystem is still unhabituated [35], while 31% of the Virunga mountain gorilla population is unhabituated [2]. The number of habituated groups in Bwindi has been increasing [35, 37, 59], and the government may decide to habituate additional gorilla groups in the future. In addition, the decision makers may try to change or disregard the current tourism regulations/guidelines (e.g., a maximum of eight tourists in one tourist group, and only one visit by one tourist group for 1 h per day per group) to increase tourism revenue. While it is too early to determine whether mountain gorillas in Bwindi were chronically stressed due to tourism, the combinations of multiple, simultaneous stressors may have detrimental effects on their immune and reproductive systems [60], which may significantly impact the survival of this species. Therefore, we recommend strictly following the tourism regulations/guidelines, specifically by limiting tourist visits to a maximum of once per day and maintaining a group size of no more than eight individuals per visit. Finally, we strongly encourage further long-term and large-scale studies targeting a larger number of gorilla groups with more manpower and higher sampling frequency. Such efforts are essential to obtain more detailed information on the influence of gorilla behavioral patterns, intra- and inter-group interactions, and varying levels of habituation. This would provide a more complete picture of the relationship between tourism activity and the physiological stress, helping to elucidate how these external and internal factors collectively shape the stress profiles of mountain gorillas.

## Acknowledgment

We thank the Uganda Wildlife Authority and Uganda National Council for Science and Technology for giving permission to conduct this research in Bwindi Impenetrable National Park, Uganda. We thank all ranger guides, trackers, and management staff of UWA for helping us and offering advice on conducting the fieldwork. We are grateful to the staff of Conservation Through Public Health and the people in the Buhoma community for their help during the fieldwork. We would like to express our gratitude to Dr. Itaru Ohta, Dr. Masayoshi Shigeta, Dr. Shiro Kohshima, and Dr. Miho Murayama for giving permission to import fecal samples from Uganda to Japan. We are indebted to Dr. Juichi Yamagiwa, Dr. Martha Robbins, Dr. Winnie Eckardt, and Dr. Raquel Philomena, who offered helpful advice and constructive comments on the research plans. We thank Dr. Shuichi Oyama, Dr. Hiroki Sato, and two anonymous reviewers for providing helpful feedback to the early version of this manuscript. Finally, we would like to thank the gorillas in Bwindi.

## Data availability statement

The data and source code are available from https://github.com/ryoma-otsuka/mg-fgcms.

## Author contributions

RO conceived and directed the study, performed the method design, led the data collection in the field, conducted laboratory work, data analysis, software implementation, data visualisation, and wrote and revised the manuscript. GKZ supported the data collection in the field and contributed to paper writing. GY supervised RO in developing the research design and contributed to paper writing. KK supervised RO for the fecal sample collection methods and laboratory work and contributed to paper writing.

## Supporting Information

### Supplementary Experiments

We conducted the following experiments to better understand the limitations of the field-friendly methods used in this study and to provide practical recommendations for future studies.

### Experiment S1

#### The amount and ratio of feces to ethanol used for hormone extraction

In the previous studies in Virunga [25, 26], the authors extracted 0.50 g of feces in 5.0 mL of 90% ethanol. However, we extracted 1.00 g of feces to reduce measurement errors when using a portable scale and used 8.0 mL of 90% ethanol to minimize ethanol consumption. The ratio of sample wet weight to 90% ethanol volume used in this study is the same as those used in Experiments 2–4 in [44]. To test whether the amount and ratio of feces to ethanol ratio for extraction has an effect on extraction efficacy, we compared the two methods: extracting 1.00 g of feces in 8.0 mL of 90% ethanol, and extracting 0.50 g of feces in 5.0 mL of 90% ethanol. Using 31 fecal samples, we extracted each sample in two ways and measured the fGCMs concentrations. We used a Bayesian model to estimate the mean difference among each pair of samples using a normal distribution. The model structure and priors are as below.

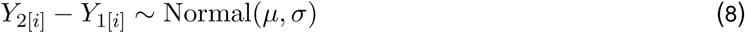

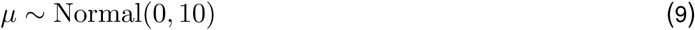

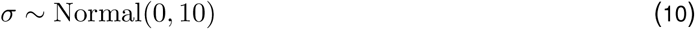

*Y*_2[_*_i_*_]_ and *Y*_1[_*_i_*_]_ represent log-fGCMs concentrations of sample i, extracted in the two different methods. The model assumes that the difference values follow a normal distribution with a mean of *µ* and a standard deviation of *σ*. For MCMC sampling, we used four chains, 4,000 iterations with 2,000 warm-up, and samples were not thinned out. We confirmed the convergence of MCMC sampling in the same way we have described in the main text. The same model, including the priors and settings, was used for Experiments S2 and S3.

In 21 out of 31 fecal samples (70.09%), log-fGCMs (and fGCMs) concentrations were higher when extracting 0.50 g of feces in 5.0 mL of 90% ethanol than extracting 1.00 g of feces in 8.0 mL of 90% ethanol, while we observed an inverse relationship in the other nine out of 31 fecal samples (29.03%) (Figure S1ad). The median (97% BCI) values of estimated posterior distributions of parameters *µ* and *σ* were 0.228 (0.073–0.383) and 0.398 (0.305–0.546), respectively. Although there was a wide uncertainty, the results suggest that extracting 0.50 g of feces in 5.0 mL of 90% ethanol would generally result in higher log-fGCMs (and fGCMs) concentrations than extracting 1.00 g of feces in 8.0 mL of 90% ethanol. Therefore, it is recommended that future studies use 0.50 g of feces and 5.0 mL of 90% ethanol to efficiently extract fGCMs from the fecal samples of mountain gorillas following the previous studies in Virunga [25, 26].

### Experiment S2

#### Extraction without a filter versus extraction using a filter

A paper filter was used for hormone extraction in Eckardt et al., [25], while a paper filter was not used in Shutt et al. [27, 44]. We did not use a paper filter for the samples used in the main analysis. To test whether the use of a paper filter affects extraction efficacy, we extracted hormones from 34 fecal samples in two different ways: using a filter (Filter Paper No. 1, 110mm, ADVANTEC), and without the use of a paper filter. We used the Bayesian model described in Experiment S1 to estimate the mean difference between each pair of samples.

In 23 out of 34 fecal samples (67.65%), log-fGCMs (and fGCMs) concentrations were higher when using a paper filter than without using it, while we observed an inverse relationship in the other 11 out of 34 fecal samples (32.35%) (Figure S1be). The median (97% BCI) values of estimated posterior distributions of parameters *µ* and *σ* were 0.173 (0.059–0.290) and 0.301 (0.232–0.400), respectively. Although there was a wide uncertainty, similar to Experiment S1, the results suggest that the use of a filter would generally result in higher log-fGCMs (and fGCMs) concentrations compared to not using a filter. Removing small particles using a filter may enhance the hormone extraction efficacy. Therefore, it is recommended that future studies use a filter to efficiently extract fGCMs from the fecal samples of mountain gorillas following Eckardt et al., [25].

### Experiment S3

#### Trail versus nest samples

Sampling from nests is much easier and less invasive than sampling from trails that require the continuous tracking of gorillas. Although gorillas defecate several times a day, it is challenging to collect fecal samples from target gorillas in a limited time (maximum 4 h of observation per day). To test if we can use nest samples along with fresh trail samples in the data analysis, we collected fecal samples from trails and the previous night nests of the three identified silverback gorillas. In total, 40 pairs of trail and nest samples were obtained (24, 11, and 5 pairs from the silverbacks Kabukojo, Maraya, and Kanyonyi, respectively).

Each of the three groups included only one silverback at the time of the fieldwork; therefore, nest samples could easily be identified by the dung size and the silvery hairs left in the nest (e.g., [61]). We used the Bayesian model described in Experiment S1 to estimate the mean difference among each pair of samples.

In 21 out of 40 fecal samples (52.50%), log-fGCMs (and fGCMs) concentrations were higher in nest samples than trail samples, while we observed an inverse relationship in the other 19 out of 40 fecal samples (47.50%) (Figure S1cf). The median (97% BCI) values of posterior distributions of parameters *µ* and *σ* were −0.005 (−0.129–0.111) and 0.334 (0.264–0.433), respectively. As such, there was no consistent relationship between log-fGCMs concentrations in the trail and nest samples and there was a wide uncertainty. A previous study in Bwindi [23] suggested that nest samples may be utilized in fGCMs analysis along with trail samples based on the results of Spearman’s rank correlation analysis and the null hypothesis testing (Wilcoxon signed rank test). When we used the same data analysis procedure as [23] for comparison, the results also showed a high correlation between trail and nest samples (Spearman’s rank correlation: rho = 0.755, p *<* 0.001), and that there was no significant difference between them (Wilcoxon signed rank test: V = 380, p = 0.695). However, it is important to note that correlation is based on ranks, and the results of the null hypothesis testing do not mean that there is no difference between the two groups. As discussed in [23], it is likely that gorillas defecate in their nests during early morning, just before they start moving (leaving their nests). However, nest samples will be left in the forest environment for several hours (after defecation until the sample collection), and this may affect the fGCMs concentrations. We further discuss this point in Experiment S4 below. Besides, the individual identification is often required to include a random effect (intercept or slope) in statistical models based on the assumption that each gorilla has a different fGCMs baseline or response pattern. If individuals can be reliably identified from nests or dung size, sampling from nests can be useful. However, unlike the western lowland gorilla population, such a situation is not very common in the mountain gorilla populations in which the percentage of multi-male groups can range from 8% to 53% [62]. Therefore, we conservatively interpreted the results from Experiment S3 and excluded nest samples in our data analysis. We also recommend that future studies carefully consider whether to use samples obtained from nests in their data analyses.

### Experiment S4

#### Post defecation changes in fGCMs concentrations

To see if fGCMs concentrations changed after defecation, we conducted the following field experiment on 29th May 2018 and 3rd October 2018 using five fecal samples, respectively. Following the early morning collection of fecal samples in the forest, we placed the collected fecal samples in an ice bag and immediately returned to the camp. We conducted the first hormone extraction for each sample at approximately 2 h after defecation. Then, we placed each fecal sample on a leaf and left it in a forest-like environment within the camp. We performed hormone extraction from each sample every 2 h up to 14 h after the defecation time, resulting in seven extracted samples (2, 4, 6, 8, 10, 12, and 14 h) for each of the 10 fecal samples. On 3rd October 2018, samples were occasionally exposed to light rainfall after extraction at 4 h up to extraction at 10 h; this did not occur on 29 May 2018. We visualised the temporal changes in fGCMs (and log-fGCMs) concentrations in Figure S2.

The fGCMs (and log-fGCMs) concentrations fluctuated from 2 h (the initial extraction in this experiment) up to 14 h after defecation; however, there was no consistent trend (increase or decrease) (Figure S2). Although fGCMs (log-fGCMs) concentrations were relatively stable in eight of ten fecal samples, there was a drastic decrease in two fecal samples.

A previous study concluded that fecal samples remain viable for corticoid analysis up to 60 h after defecation; however, in practical applications, fecal samples collected within 12 h post-defecation (i.e., samples from previous night nests) should be used [23]. However, Figure 1A in [23] showed a roughly increasing trend in fGCMs concentrations up to 60 h, while it also showed an inverse trend at the same time (e.g., see the decreasing trend in the sample from the Silverback, indicated as SB, from 0 to 24 h). The results from this study indicate that measured fGCMs concentrations drastically decreased in two of 10 samples between 2 and 4 h after defecation. The reason behind the drastic change is unclear, but it may be due to measurement error or the influence of rain on 3rd October 2018. Although we did not observe such drastic changes in other samples, at times, measured fGCMs concentrations in some samples had either doubled or been reduced by half. Therefore, we conservatively interpreted that the results of the previous study [23] and our Experiment S4; we concluded that fGCMs concentrations would change after defecation, and this trend may not be consistent, affected by various factors.

There are three possible causes for this post-defecation change in the fGCMs concentrations of mountain gorillas in Bwindi. First, contamination and measurement errors may have affected fGCMs concentrations, and it is necessary to take all possible measures to reduce contamination and other measurement errors. Second, ultraviolet light may have also affected fGCMs concentrations [63], and ultraviolet light may be avoided by placing collected samples into a container that prevents light exposure. Third, the activity of bacteria in feces may have influenced fGCMs concentrations [63], and bacterial activity may be reduced under low temperatures. Therefore, it is strongly recommended that future research use a cooling box or ice bag with ice packs that can maintain collected fecal samples at a low temperature until they can be stored in the fridge in the camp, as we did in this study. This would also prevent fecal samples from being exposed to direct sunlight during sample transportation. Yet, it is not unclear whether or how much post-defecation changes can be minimized using these techniques. We recommend that future studies investigate these effects and develop methods that are both robust and easy for field researchers to implement.

## Supplementary Figures

**Figure S1.**
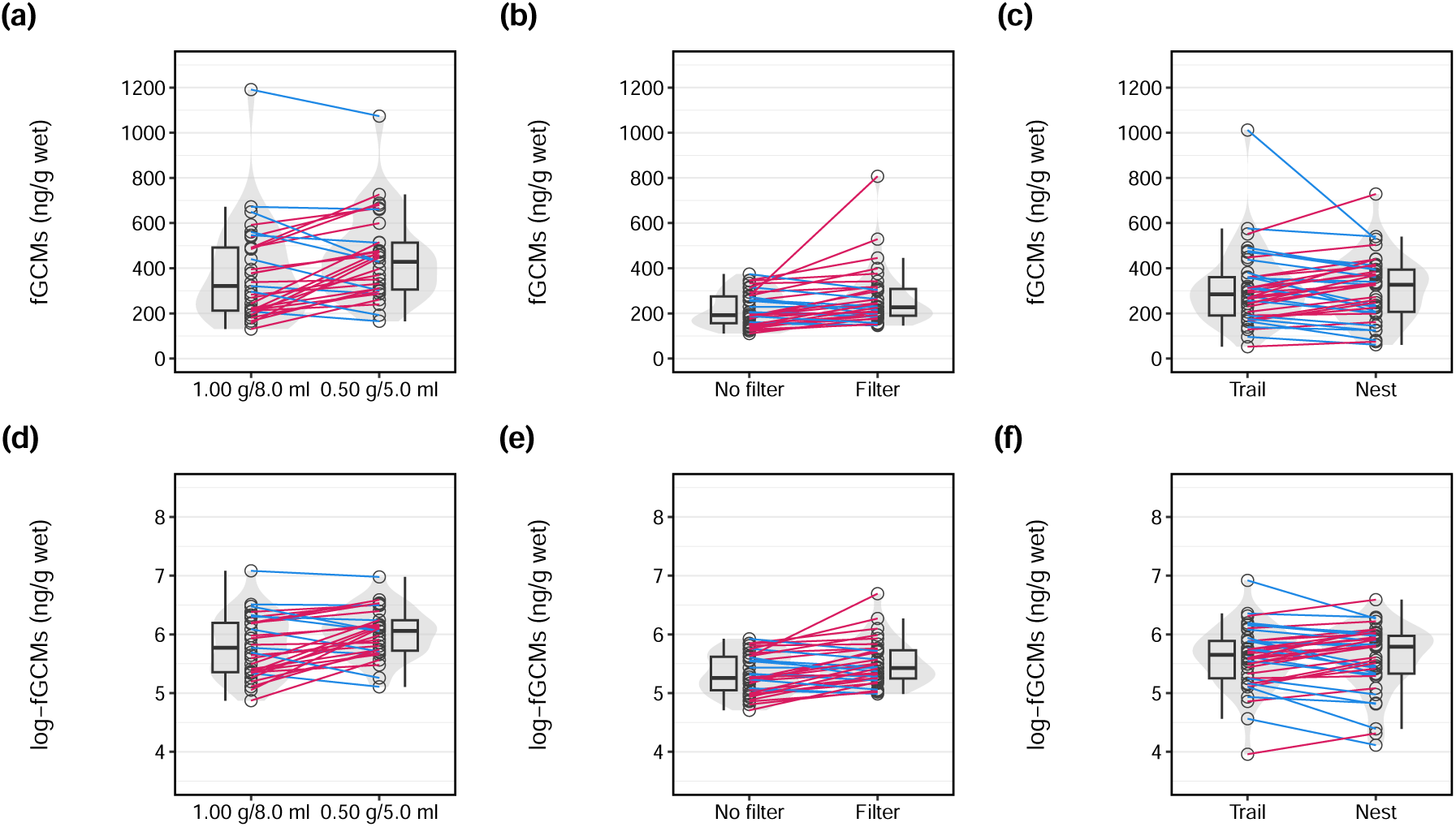
Comparison of fGCMs concentrations between (a) two different extraction methods: 1.00 g of feces in 8.0 mL of 90% ethanol versus 0.50 g of feces in 5.0 mL of 90% ethanol (31 pairs of samples), (b) two different extraction methods: using no filter versus using a filter (34 pairs of samples), and (c) trail and nest samples collected on the same day from the same silverback gorilla (40 pairs of samples from three silverback gorillas). The plots (d), (e), and (f) are the log-transformed versions of (a), (b), and (c), respectively.

**Figure S2.**
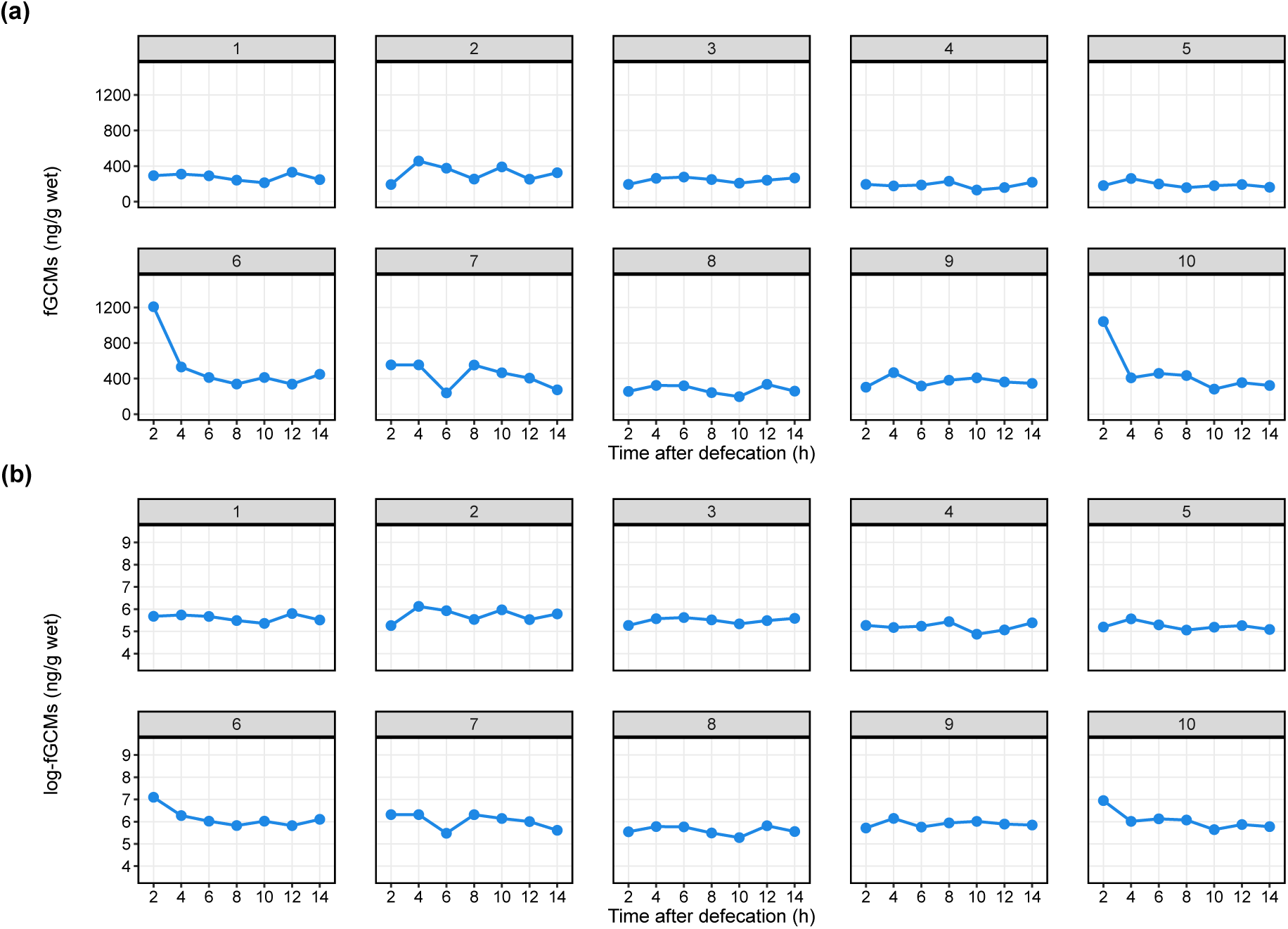
Post defecation changes in fGCMs concentrations. (a) Post defecation changes in fGCMs concentrations from 2 up to 14 h after defecation (N = 10). The fecal samples 1–5 were extracted on 29 May 2018, and the samples 6–10 were extracted on 3 October 2018. The plot (b) is the log-transformed version of (a).

**Figure S3.**
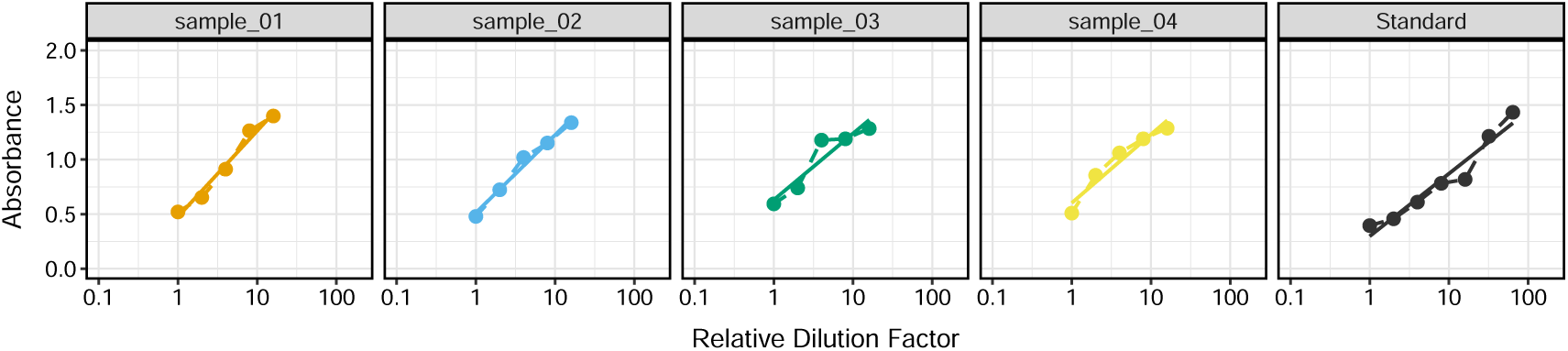
Parallelism checks of enzyme immunoassay. Each circle indicates the mean absorbance of duplicated samples. The dotted lines simply connect each data point, while the solid lines are linear regression lines.

**Figure S4.**
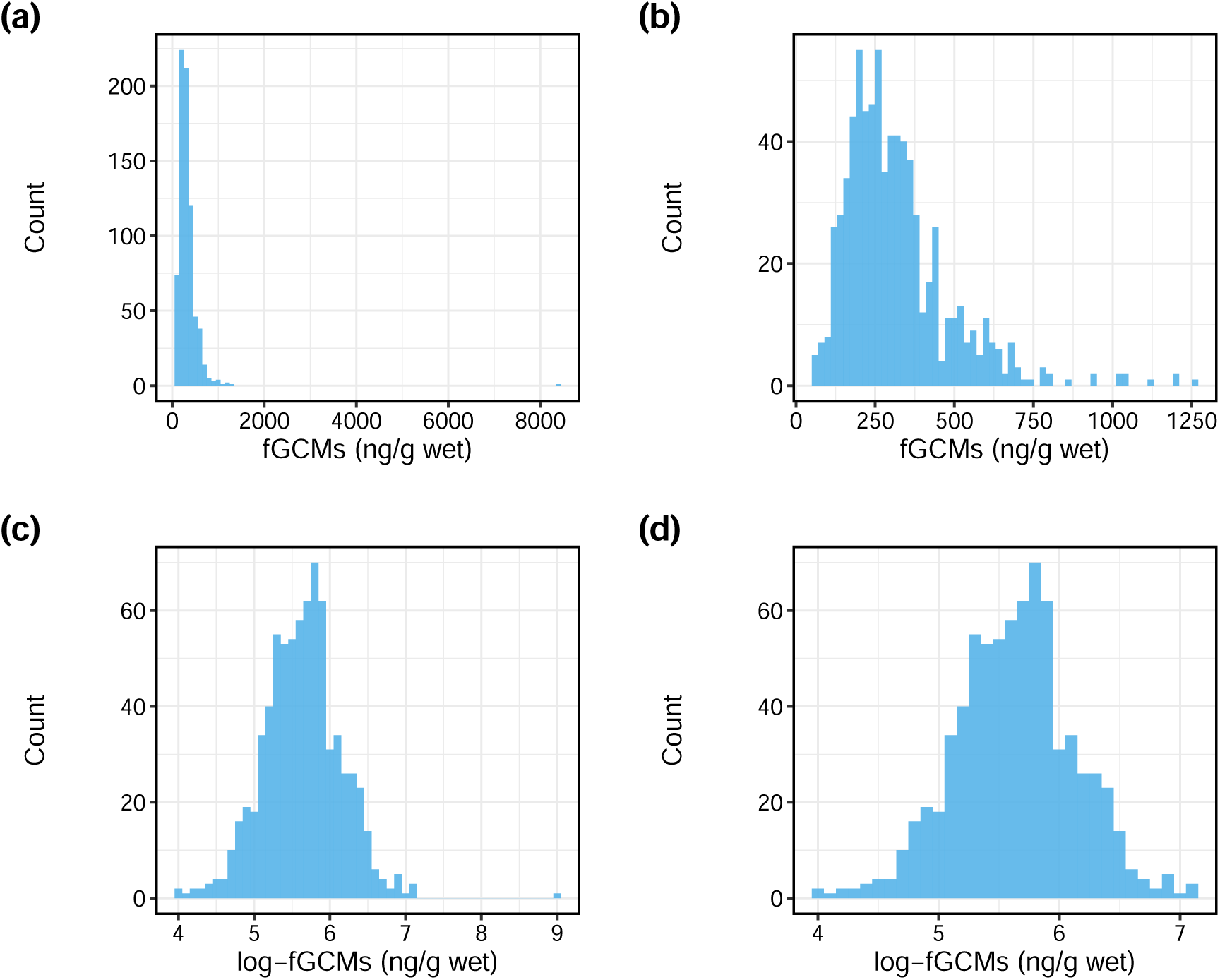
Distribution of fGCMs (a, b) and log-transformed fGCMs concentrations (c, d). An extreme value (*>*8,000 in fGCMs) shown in (a, c) was removed in (b, d).

